# Pre-existing heterogeneity facilitates development of heteroresistance upon gene acquisition

**DOI:** 10.1101/2023.07.24.550411

**Authors:** Siddharth Jaggavarapu, David A. Hufnagel, David S. Weiss

## Abstract

Antibiotic resistance causes 1.27 million global deaths annually and is predicted to worsen. Heteroresistance is a form of resistance in which only a minor and unstable subpopulation of cells of a bacterial isolate are resistant to a given antibiotic, and are therefore often undetected by clinical diagnostics. These infrequent and undetected resistant cells can be selected during antibiotic therapy, expand in number, and cause unexplained treatment failures. A major question is how heteroresistance evolves. Here, studying the antibiotic fosfomycin, we report that heteroresistance can develop from a pre-existing state of phenotypic heterogeneity in which an isolate harbors a subpopulation with increased minimum inhibitory concentration (MIC), but below the clinical resistance breakpoint. We call this phenomenon heterosusceptibility and demonstrate that acquisition of a resistance gene, *fosA*, increases the MIC of the subpopulation beyond the breakpoint, making the isolate heteroresistant. Conversely, deletion of *fosA* from a heteroresistant isolate led to reduction of the MIC of the resistant subpopulation without a loss of heterogeneity, thus generating heterosusceptibility. A survey of 103 carbapenem-resistant Enterobacterales (CRE) revealed that the *Escherichia sp*. isolates lacked the *fosA* gene and uniformly exhibited fosfomycin heterosusceptibility, whereas the *Klebsiella* and *Enterobacter* encoded the *fosA* gene and were almost exclusively heteroresistant. Furthermore, some isolates exhibited heterosusceptibility to other antibiotics, demonstrating that this is a widespread phenomenon. These results highlight a mechanism for the evolution of heteroresistance and suggest that surveillance for heterosusceptibility may facilitate the prediction of impending heteroresistance before it evolves.

## Introduction

Antibiotic resistance is estimated to cause 1.27 million deaths per year worldwide and is thus one of the greatest medical challenges of the 21^st^ century (1). Without new drugs, it is predicted that antibiotic resistant infections will cause 10 million annual deaths worldwide by the year 2050 (surpassing cancer) and by that time add $100 trillion to the world’s healthcare costs (2). Drug-resistant infections are poised to negate many advances of modern medicine which rely on antibiotics including transplants, chemotherapy, and survival of extremely premature infants, and may even preclude routine or elective surgeries.

One understudied form of antibiotic resistance is heteroresistance (HR), in which a minor subpopulation of resistant bacterial cells co-exists with a susceptible majority population. Even when as infrequent as 1 in 1 million, the resistant cells rapidly expand in the presence of a given antibiotic and thus can cause treatment failure (3, 4). Due to the low frequency of resistant cells, heteroresistance is often undetected by clinical diagnostics (4, 5).

A major outstanding question is how bacteria evolve heteroresistance. A standard view of the evolution of conventional, homogenous antibiotic resistance is that it occurs following acquisition of a resistance gene or mutation, and that all cells in the population have a uniformly elevated minimum inhibitory concentration (MIC). Similarly, lack of the resistance gene or mutation in the parental, susceptible strain leads to all cells being equally susceptible to a given antibiotic. However, this model does not account for the occurrence of population heterogeneity in either parental susceptible strains or evolved resistant strains.

Here, we reveal that bacterial isolates can harbor subpopulations with differing MICs but which all survive only at low concentrations of antibiotic below the clinical breakpoint. We use the term heterosusceptibility to describe this phenomenon, in contrast to heteroresistance in which a subpopulation survives above the clinical breakpoint. We identify clinical *E. coli* isolates exhibiting heterosusceptibility to fosfomycin, an antibiotic that inhibits the MurA peptidoglycan synthesis protein and thus interferes with bacterial cell wall synthesis (6). Interestingly, a fosfomycin heterosusceptible *E. coli* isolate became heteroresistant upon introduction of a fosfomycin resistance gene (*fosA*) whose gene product covalently inactivates the drug (7). These data demonstrate that acquisition of a resistance gene can increase the MIC of a heterogeneously fosfomycin-susceptible strain to generate heteroresistance. These findings significantly expand our understanding of the evolution of heteroresistance and have important implications for the future study and epidemiological monitoring of antibiotic resistance.

## Results

We previously reported that a strain of *Enterobacter cloacae*, Mu208, exhibits heteroresistance to fosfomycin (Figure 1A) (8). This phenotype was detected using population analysis profile (PAP), the gold standard method for detecting heteroresistance (2). In a PAP assay, bacteria are plated on agar with or without increasing antibiotic concentrations and the proportion of resistant cells is calculated. From the Mu208 PAP data, we observed distinct subpopulations of cells and subsequently determined the MIC of the susceptible subpopulation (MIC-S), the lowest fosfomycin concentration at which >90% of the cells were killed. The MIC-S was 256 μg/ml fosfomycin, which is also the clinical resistance breakpoint, the concentration at and above which bacterial survival is associated with clinical treatment failure (Figure 1A and Table 1). In contrast, the MIC of the resistant subpopulation (MIC-R), the fosfomycin concentration at which all the cells would be killed, was >1,024 μg/ml, above the clinical breakpoint, demonstrating heteroresistance (Figure 1A and Table 1).

**Table 1.** MICs of distinct subpopulations of cells in heterosusceptible and heteroresistant strains.

| Isolate | Classification | MIC-S | MIC-R |
| --- | --- | --- | --- |
| Mu208-WT | Heteroresistant | 256 | >1024<sup>b</sup> |
| Mu208Δ*fosA* | Heterosusceptible | 4 | 128 |
| Mu208Δ*fosA* att::*fosA* | Heteroresistant | 256 | >1024<sup>b</sup> |
| Mu582-WT | Heterosusceptible | <0.5<sup>a</sup> | 16 |
| Mu582 att::*fosA* | Heteroresistant | 1 | 1024 |

Fosfomycin population analysis profile (PAP) was performed on each of the indicated strains which were subsequently classified as either heterosusceptible or heteroresistant. The MIC of the most susceptible subpopulation (MIC-S) and the subpopulation with the highest MIC (MIC-R) was determined for each strain.
<sup>a</sup>MIC was lower than the lowest concentration tested by PAP
<sup>b</sup>MIC was higher than the highest concentration tested by PAP

**Figure 1.**
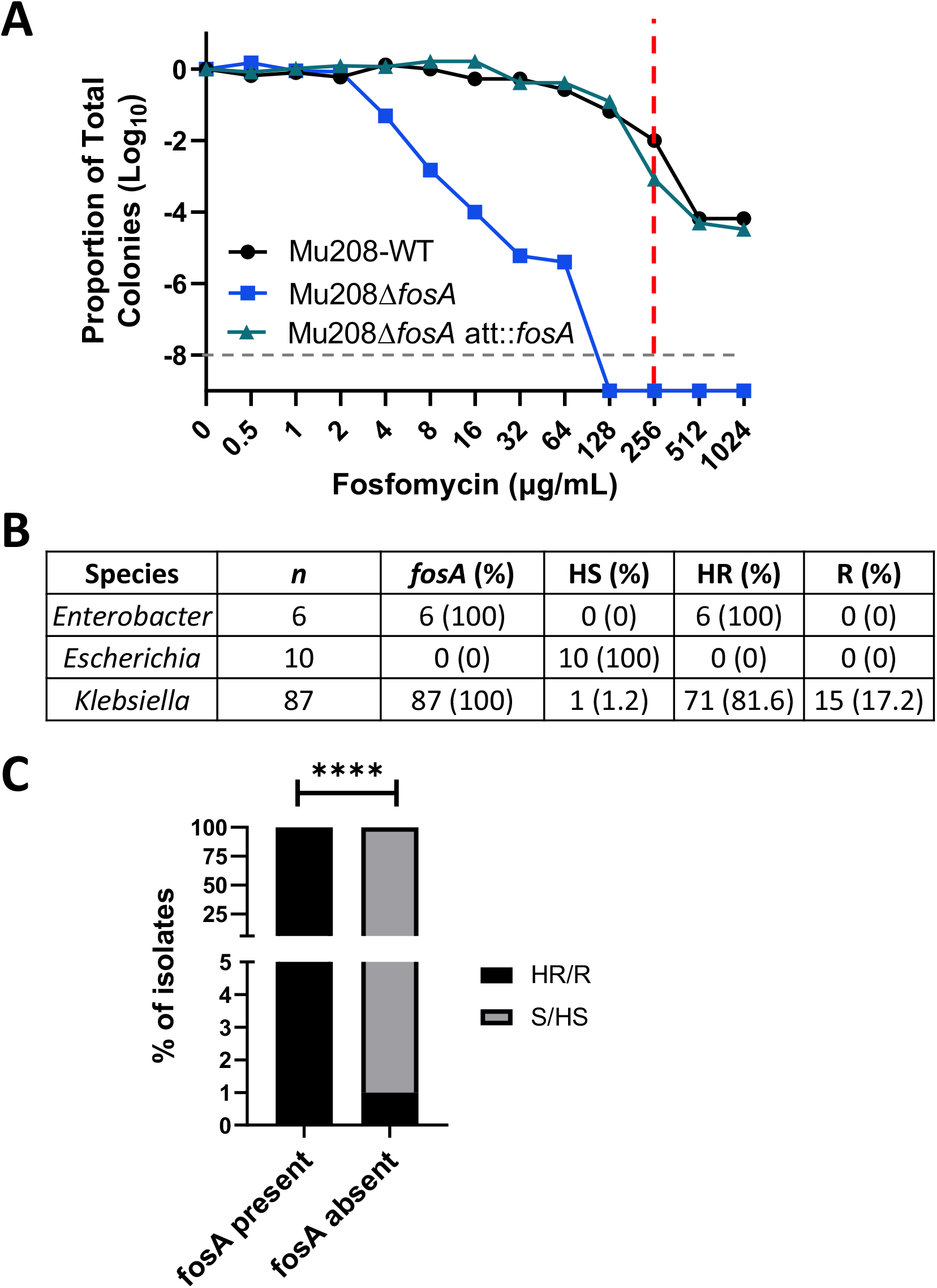
Presence of the *fosA* gene is associated with fosfomycin heteroresistance. (A) Population analysis profile (PAP) of the *Enterobacter cloacae* strain WT (Mu208-WT), *fosA* deletion (Mu208D*fosA*) and the complemented strain (Mu208D*fosA* att::*fosA*) plated on the indicated concentrations of fosfomycin. The proportion of total colonies was calculated relative to the total CFUs on the antibiotic-free plates. The dashed red-line indicates the fosfomycin breakpoint and the dashed grey line is the limit of detection of the assay. (B) The genomes of 103 carbapenem-resistant Enterobacterales from the indicated species were evaluated for the presence of a *fosA* gene. The number of isolates that are heterosusceptible (HS), heteroresistant (HR), or resistant (R) according to PAP analysis is indicated. (C) The presence of a *fosA* gene is associated with fosfomycin heteroresistance. The P-value (\*\*\*\**P* <0.0001) was obtained by comparing the data using Fisher’s exact test.

The genome of Mu208 contains a fosfomycin resistance gene, *fosA*, and we investigated whether this gene contributed to the heteroresistance phenotype. A *fosA* deletion mutant (Mu208D*fosA*) was susceptible to fosfomycin as assayed by PAP; no cells survived at the clinical breakpoint of 256 ug/ml (Figure 1A). However, a close examination of the PAP curve of the Mu208D*fosA* strain revealed the presence of a subpopulation of cells surviving up to 64 μg/ml, in contrast to the majority population which was killed at 4 μg/ml (Figure 1A). Therefore, Mu208D*fosA* exhibits heterogeneity similar to the parental Mu208 strain, but with all cells surviving only at lower fosfomycin concentrations below the clinical breakpoint, which we refer to here as heterosusceptibility. Importantly, these data indicated that the heterogeneity phenotype was independent of *fosA* (Figure 1A and Table 1). Rather, the presence of *fosA* relatively uniformly increased the MIC of the heterogeneous population by 16 to 32-fold (Figure. 1A and Table 1). Complementation of Mu208D*fosA* with a single chromosomal copy of the *fosA* gene (creating strain Mu208D*fosA* att::*fosA*) led to restoration of heteroresistance, as shown by a resulting PAP curve similar to that of the parental Mu208 strain and with a subpopulation surviving at 1,024 μg/ml, above the resistance breakpoint (Figure 1A). These data demonstrate that while loss of a resistance gene from a fosfomycin heteroresistant isolate leads to a reduction in the concentration up to which the strain can survive, the heterogeneity of the strain is retained.

Heteroresistance is a form of phenotypic heterogeneity in which the frequency of the resistant subpopulation is unstable. Accordingly, in Mu208 and Mu208D*fosA* att::*fosA*, the resistant subpopulation was enriched and became predominant after growth in the breakpoint concentration of fosfomycin, and subsequently reverted to the baseline frequency after growth in drug-free media (Supplemental Figure 1). We next investigated whether the Mu208D*fosA* deletion strain exhibited similar phenotypic instability. At a low concentration of fosfomycin (4 μg/ml) below the clinical breakpoint and at which Mu208D*fosA* could survive, the subpopulation with a higher MIC than the majority population was enriched (Supplemental Figure 2). After subsequent growth in media without fosfomycin, the frequency of this subpopulation reverted to baseline. Therefore, heterosusceptibility is characterized by phenotypic instability, similar to heteroresistance, but at sub-breakpoint concentrations.

Since *fosA* can have such a profound impact on heteroresistance to fosfomycin, we investigated the presence of *fosA* among a collection of 103 carbapenem resistant isolates of three Enterobacterales species from Georgia, USA. Whole genome sequencing data revealed that all the *Klebsiella* and *Enterobacter* species encoded *fosA*, while none of the 10 *Escherichia* encoded the gene (Figure 1B). These data are in line with those observed by Ito et al (9), who analyzed the genomes of 18,130 Gram-negative bacterial species and found that only 4.6% of the *E. coli* genomes encoded a *fosA* gene, whereas >95% of *Klebsiella* and *Enterobacter* species encoded a *fosA* gene.

We next tested whether the presence of the *fosA* gene correlated with fosfomycin susceptibility status. Indeed, almost all of the *Enterobacter* and *Klebsiella* isolates tested were fosfomycin heteroresistant, while none of the *Escherichia* isolates tested exhibited this phenotype (Figure 1B). However, all the *Escherichia* isolates did exhibit heterogeneity below the resistance breakpoint and were thus classified heterosusceptible to fosfomycin by our PAP analyses (Figure 1B). Taken together, these data indicate that there is a strong correlation between *fosA* and fosfomycin heteroresistance, and likewise between the lack of *fosA* and fosfomycin heterosusceptibility (p<0.0001) (Figure 1C).

Since the *Escherichia* in our isolate collection lacked *fosA* and were uniformly heterosusceptible to fosfomycin, we questioned whether *Escherichia* could become heteroresistant to fosfomycin upon acquisition of *fosA*. We therefore cloned a single copy of the *fosA* gene into the *att-*site of the genome of a fosfomycin-heterosusceptible *E. coli* strain (Mu582). PAP analysis revealed that acquisition of *fosA* significantly increased the MIC of the resistant subpopulation in the *fosA*-encoding isolate (Mu582 att::*fosA*) above the clinical breakpoint, such that this strain became heteroresistant (Figure 2). Furthermore, the resistant subpopulation was unstable; it was enriched when Mu582 att::*fosA* was grown in the presence of fosfomycin, but its frequency subsequently decreased upon passage in drug-free media (Supplemental Figure 1). These findings indicate that heterosusceptible isolates with heterogeneous subpopulations that all survive at differing drug concentrations below the clinical breakpoint can gain a resistance gene, increase their MIC, and become heteroresistant.

**Figure 2.**
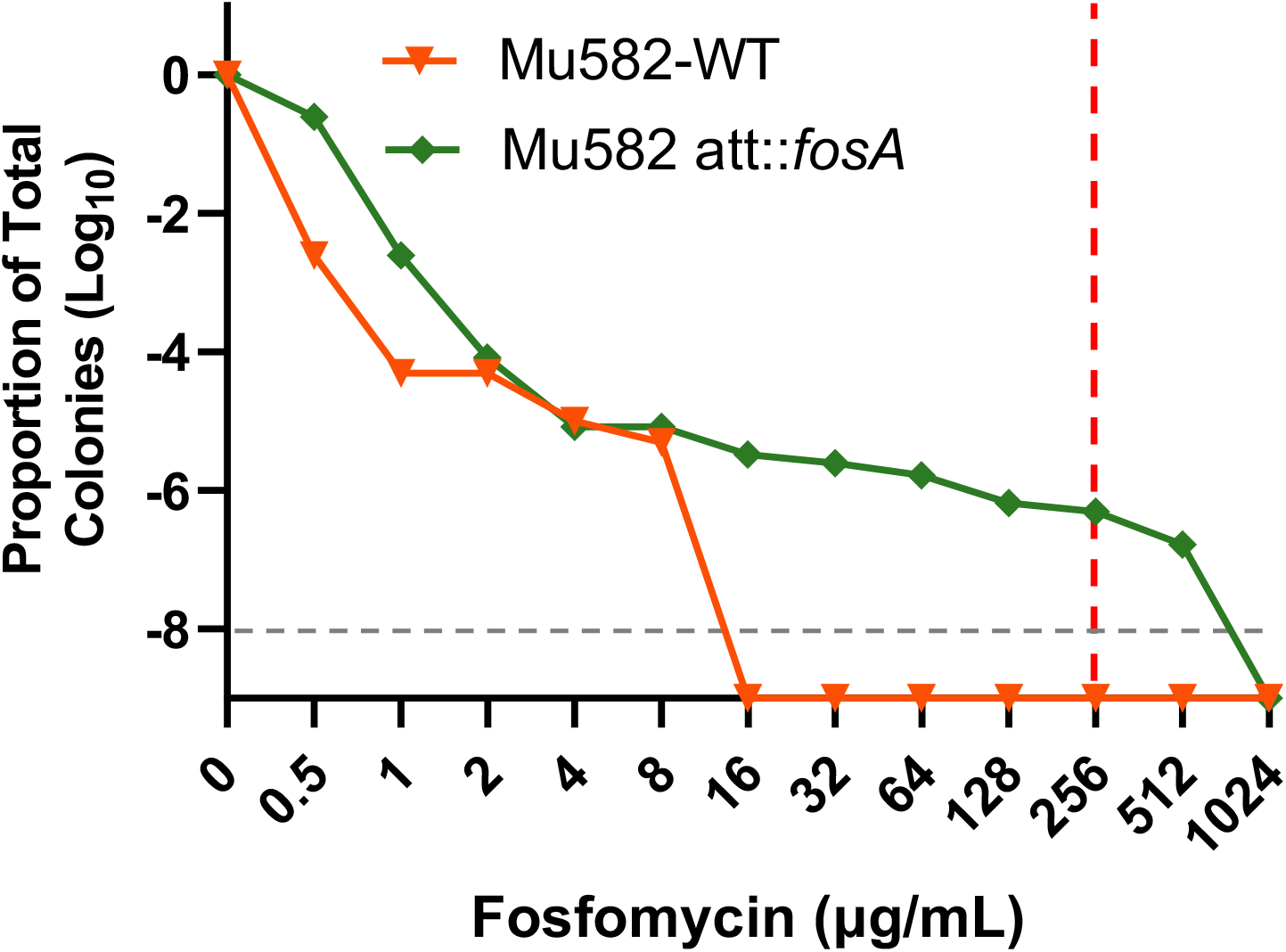
Acquisition of *fosA* converts a fosfomycin heterosusceptible isolate to heteroresistant. PAPs of the fosfomycin heterosusceptible *E. coli* strain Mu582 and the *fosA* complemented strain (Mu582 att::*fosA*) plated on fosfomycin plates at indicated concentrations. The proportion of total colonies was calculated relative to the number of colonies on the antibiotic-free plates. The dashed red line indicates the fosfomycin breakpoint and the dashed grey line is the limit of detection of the assay.

Heteroresistance has been shown to cause antibiotic treatment failure using *in vivo* infection models (5, 8). We thus investigated the impact of heterosusceptibility on fosfomycin treatment outcome using the *Galleria mellonella* waxworm infection model. The heteroresistant Mu208 strain killed 85% of infected worms in the absence of antibiotic treatment (Supplemental Figure 3A). Fosfomycin treatment failed to rescue the infected worms, consistent with heteroresistance causing treatment failure (Supplemental Figure 3A). Similarly, the heterosusceptible Mu208Δ *fosA* strain killed 95% of infected worms. However, fosfomycin treatment was effective as it rescued 80% of the worms infected with Mu208D*fosA* (Supplemental Figure 3B). Similar results were observed with *E. coli*, where fosfomycin treatment only rescued the worms infected with the parental heterosusceptible Mu582 strain, but not with the heteroresistant Mu582 att::*fosA* strain (Supplemental Figures 3C, D). Taken together, these data demonstrate that in contrast to heteroresistance, heterosusceptibility is not associated with fosfomycin treatment failure in the waxworm model, but rather can be a precursor to heteroresistance.

The results presented here, highlighting the phenomenon of fosfomycin heterosusceptibility, led us to investigate whether heterosusceptibility exists for other classes of antibiotics. Among our collection of 103 CRE, we surveyed all instances in which isolates were previously classified susceptible to one of a panel of 17 antibiotics by PAP (8). We observed heterosusceptiblity for almost every antibiotic tested, including beta-lactams, polymyxins, quinolones, sulfonamides, tetracyclines, and ranging from 8% (ceftazidime-avibactam) to 100% (fosfomycin) of isolates previously classified susceptible (Supplemental Table 1). These data demonstrate that heterosusceptibility is a phenomenon which occurs for many antibiotics from diverse classes.

## Discussion

The mechanisms by which bacteria evolve heteroresistance have not been well understood. In this study, we identify a precursor of antibiotic heteroresistance which we term heterosusceptibility. Heterosusceptibility occurs when there is heterogeneity in the MIC of a subpopulation of cells of a given isolate and to a given antibiotic, but in contrast to heteroresistance, the MICs of all cells in the population are below the clinical resistance breakpoint (Figure 1A and Supplemental Figure 4). Importantly, as with heteroresistance, in heterosusceptibility the frequency of the subpopulation with increased MIC is unstable and increases when the isolate is exposed to a given antibiotic, but reverts to baseline in the absence of drug (Supplemental Figure 2).

We show that resistance gene acquisition can facilitate the transition from heterosusceptibility to heteroresistance (Figure 2). All the *Escherichia* isolates in our collection of 103 CRE lacked the *fosA* gene and were heterosusceptible to fosfomycin (Figure 1B). In contrast, all the *Klebsiella* and *Enterobacter* isolates encoded the *fosA* gene, and all but one *Klebsiella* isolate were heteroresistant to fosfomycin (Figure 1B). Interestingly, when we engineered a single copy of the *fosA* gene into the genome of an *E. coli* isolate, the strain shifted from fosfomycin heterosusceptible to heteroresistant (Figure 2). Conversely, deletion of *fosA* from an *Enterobacter cloacae* isolate which was heteroresistant to fosfomycin rendered the strain heterosusceptible (Figure 1A).

Interestingly, it is generally assumed that acquisition of a resistance gene causes all cells in a population to become resistant. In contrast, the data presented here show this is not always the case and that a heterogeneous response may occur. Therefore, heteroresistance may explain at least some cases in which acquisition of a known resistance gene does not lead to resistance to a corresponding antibiotic when assayed by standard antimicrobial susceptibility testing (AST), since these tests often lack the sensitivity to detect heteroresistance.

Previously, one study demonstrated that continuous exposure to sub-MIC concentrations of colistin led to the evolution of heteroresistance via amplification (increased gene copy number) of genes involved in colistin resistance (10). It will be of great interest in the future to identify additional ways in which heteroresistance can evolve. In addition, it will be critical to elucidate the mechanistic basis of heterosusceptibility and how heterogeneity is generated in the absence of a given resistance gene, for example in fosfomycin heterosusceptibility in the absence of *fosA*.

We suggest that epidemiological surveillance networks should not only consider screening for heteroresistance, but also for heterosusceptibility, as this could be an indicator of the impending, future development of heteroresistance. Data on susceptibility, heterosusceptibility, heteroresistance, and resistance would provide epidemiologists, clinical microbiologists, and clinicians with the most complete picture of the overall antibiotic resistance landscape, potentially informing antibiotic stewardship and clinical practices.

## Materials and Methods

### Bacterial isolates

The CRE isolates used in this study were collected between 2013 and 2015 by the Georgia Emerging Infections Program’s Multisite Gram-negative Surveillance Initiative, as described previously (11). The Mu208Δ *fosA* strain was created by deleting the *fosA* gene from the Mu208 genome using the lambda red recombination method as described elsewhere (12). Briefly, the *fosA* gene from Mu208 along with 200bp of upstream region was amplified by PCR with primers (P1 and P2) containing sequences complementary to the pCD13pSK vector. pCD13pSK was amplified with inverse PCR primers (P3 and P4) that excluded the multiple cloning site and the T7 promoter. The *fosA* PCR product was cloned into the inverse PCR product from pCD13pSK using NEbuilder Hifi assembly (New England Biolabs, Ipswich, MA). Both *fosA* complemented derivatives of Mu208Δ *fosA* and Mu582 were created by incorporating the *fosA* gene along with a 200bp upstream region from the Mu208 genome into the *att* site of each genome (12).

### Bacterial culture

Bacteria were streaked onto Mueller Hinton (MH) agar (BD Biosciences, NJ) plates from frozen glycerol stocks. Overnight cultures were grown from single colonies in MH broth at 37°C with shaking at 250 rpm. Colony forming units (CFU) were counted by plating serial dilutions in PBS on MH agar plates. The CFU were counted at the lowest distinguishable dilution.

### Population analysis profile (PAP)

PAPs were conducted as described previously, with some modifications (5). Typically, MH agar plates containing 25 μg/mL glucose-6-phosphate (G6P) were made with 0, 0.01565, 0.03125, 0.0625, 0.125, 0.25, 0.5, 1, 2 and 4 times the breakpoint concentrations of the respective antibiotic. Serial dilutions of overnight cultures were plated and the CFU were enumerated after overnight incubation at 37°C. PAPs were repeated at additional lower concentrations if no growth was observed at the lowest antibiotic concentration tested. Isolates were classified heterosusceptible if there was <50% survival over 3 doubling concentrations below 1X breakpoint and no survival at 1X the breakpoint. The MICs for subpopulations were determined from the PAP data. The MIC of the susceptible subpopulation (MIC-S) was determined to be the lowest concentration of the antibiotic at which 90% of the population was killed. The MIC of the resistant subpopulation (MIC-R) was determined to be the lowest concentration of the antibiotic at which all the cells were killed. Both MICs were confirmed with at least three biological replicates.

### *fosA* gene analysis

The genome sequences of all the CRE isolates are available on NCBI. The published genomes were searched for the presence of the *fosA* gene using the Comprehensive Antibiotic REsiatCARD algorithm (13).

### Antibiotic resistance stability assay

Overnight cultures were grown from single colonies in MH broth with 25 μg/mL G6P for % resistance determination prior to exposure to antibiotics. For antibiotic treatment, overnight cultures were diluted 1:1,000 into MH broth with 128 μg/mL fosfomycin (TCI, Portland, OR) and 25 μg/mL G6P and grown at 37°C for 20 hours with shaking at 250 rpm. Antibiotic-free subcultures were performed similarly by diluting antibiotic-grown overnights 1:1,000 in drug-free MH broth with 25 μg/mL G6P. Bacterial cultures were serially diluted in PBS and plated on MH agar plates with and without 128 μg/mL fosfomycin to determine the % resistance. In addition, similar experiments were performed for strains Mu208Δ *fosA* and Mu582 with 4 μg/mL Fosfomycin.

### *Galleria* infections

Overnight cultures of bacteria were washed twice and final suspensions were prepared in sterile PBS (Lonza). *Galleria mellonella* larvae were infected with 10 μL of the respective strains with a Hamilton syringe (Hamilton Company, Reno, NV) using a 0.27-gauge needle (BD Biosciences, NJ). Inocula used were 3x10^7^ CFU for *E. cloacae* Mu208 and Mu208Δ *fosA*, and 1x10^7^ CFU for *E. coli* strains. Worms were placed in petri dishes filled with bedding for 2 hours at 37°C before fosfomycin treatments. Worms infected with the *E. cloacae* strains were treated with 40 μg of fosfomyin in 10 μL volume, and worms infected with *E. coli* strains were treated with 2 μg of fosfomycin in 10 μL. Untreated worms were injected with 10 μL of PBS. Worms were incubated at 37°C for 96 hours and scored for survival every 24 hours.

## Acknowledgements

We thank Bikash Bogati and Jacob Choby for critical reading of the manuscript. We also thank Dr. Choby for cloning advice in generating some of the mutant strains used in this paper. This work was supported by NIH grants AI158080 and P51OD011132. DSW was also supported by a Burroughs Wellcome Fund Investigators in the Pathogenesis of Infectious Diseases award.

