## Supplemental data for "Pre-existing heterogeneity facilitates development of heteroresistance upon gene acquisition"

**Supplemental Legends**

**Supplemental Table 1. Frequency of heterosusceptibility to diverse antibiotics among carbapenem-resistant Enterobacterales (CRE) from Georgia, USA.** Population analysis profile (PAP) was performed on 103 CRE for the indicated antibiotics. The number of isolates previously categorized as susceptible (“S”) is indicated for each antibiotic. In addition, the number of the susceptible isolates determined to be heterosusceptible (“HS”) in the present study is indicated. “Overall %HS” indicates the percentage of the 103 isolates that are heterosusceptible to a given antibiotic, and “Susc %HS” indicates the percentage of the isolates previously classified susceptible that are in fact heterosusceptible based on the results in this study. Ceftaz-avi, ceftazidime-avibactam; Trimeth-sulfa, trimethoprim-sulfamethoxazole.

**Supplemental Figure 1. Fosfomycin resistant subpopulations in heteroresistant isolates are phenotypically unstable.** The frequency of the fosfomycin-resistant subpopulation of the indicated strains was quantified after growth in MH broth with G6P overnight (Pre-treatment), subsequent passage in MH broth with G6P and 128 μg/mL fosfomycin (in drug) for 24 h and subsequently in MH broth with G6P and no fosfomycin for 24 hours (subculture). The % resistance was calculated compared to the CFUs on no antibiotic plates.

**Supplemental Figure 2. Heterosusceptibility is characterized by unstable, phenotypic heterogeneity.** The frequency of the subpopulation of the indicated strains with increased MIC was quantified after growth in MH broth with G6P overnight (Pre-treatment), subsequent passage in MH broth with G6P and fosfomycin ("in drug"; 4 µg/ml for Mu208Δ*fosA*, 1 µg/ml for Mu582) for 24 h and following passage in MH broth with G6P and no fosfomycin for 24 hours (subculture). The % resistance was calculated compared to the CFUs on antibiotic-free plates.

**Supplemental Figure 3. Heterosusceptibility does not mediate fosfomycin treatment failure.** *Galleria mellonella* (n = 15 per group) were infected with (A) Mu208-WT, (B) Mu208 Δ*fosA*, (C) Mu582-WT or (D) Mu582 att::*fosA* strains and then inoculated with PBS (untreated, red) or fosfomycin (treated, blue) 2 hours post-infection. Survival was monitored for 96 hours with data recorded every 24 hours. ***$P$ <0.001 and ****$P$ <0.0001, two-sided log-rank test.

**Supplemental Figure 4. Schematic of population analysis profile (PAP) classifications for distinct resistance phenotypes.** The red squares represent data from a resistant isolate, in which all the cells survive at the breakpoint concentration. Blue triangles represent data from a heteroresistant isolate, in which there is a subpopulation of cells that survive at and beyond the breakpoint. Green inverted triangles represent data from a heterosusceptible isolate, in which there is a subpopulation of cells that are killed by an antibiotic concentration lower than the breakpoint. Black circles represent data from a susceptible isolate that does not contain a resistant subpopulation and is uniformly killed off at a low antibiotic concentration.

Supplemental Table 1

| Class | Antibiotic | Total | S | HS | Overall %HS | Susc %HS |
|---|---|---|---|---|---|---|
| Phosphonic acid | Fosfomycin | 103 | 7 | 7 | 6.8 | 100 |
| Aminoglycoside | Amikacin | 103 | 34 | 9 | 8.7 | 26.5 |
| | Gentamicin | 103 | 71 | 20 | 19.4 | 28.2 |
| | Tobramycin | 103 | 8 | 1 | 1.0 | 12.5 |
| Beta-lactam | Aztreonam | 103 | 1 | 1 | 1.0 | 100 |
| | Cefepime | 103 | 1 | 1 | 1.0 | 100 |
| | Ceftazidime | 103 | 0 | 0 | 0 | 0 |
| | Ceftaz-avi | 103 | 98 | 8 | 7.8 | 8.2 |
| | Meropenem | 103 | 3 | 3 | 2.9 | 100 |
| Polymyxin | Colistin | 103 | 20 | 19 | 18.5 | 95.0 |
| Quinolone | Ciprofloxacin | 103 | 1 | 1 | 1.0 | 100 |
| Sulfonamide | Trimeth-sulfa | 103 | 22 | 14 | 13.6 | 63.6 |
| Tetracycline | Tetracycline | 103 | 43 | 6 | 5.8 | 14.0 |
| | Tigecycline | 103 | 56 | 6 | 5.8 | 10.7 |

**Supplemental Table 1. Frequency of heterosusceptibility to diverse antibiotics among carbapenem-resistant Enterobacterales (CRE) from Georgia, USA.** Population analysis profile (PAP) was performed on 103 CRE for the indicated antibiotics. The number of isolates previously categorized as susceptible ("S") is indicated for each antibiotic. In addition, the number of the susceptible isolates determined to be heterosusceptible ("HS") in the present study is indicated. "Overall %HS" indicates the percentage of the 103 isolates that are heterosusceptible to a given antibiotic, and "Susc %HS" indicates the percentage of the isolates previously classified susceptible that are in fact heterosusceptible based on the results in this study. Ceftaz-avi, ceftazidime-avibactam; Trimeth-sulfa, trimethoprim-sulfamethoxazole.

Supplemental Figure 1

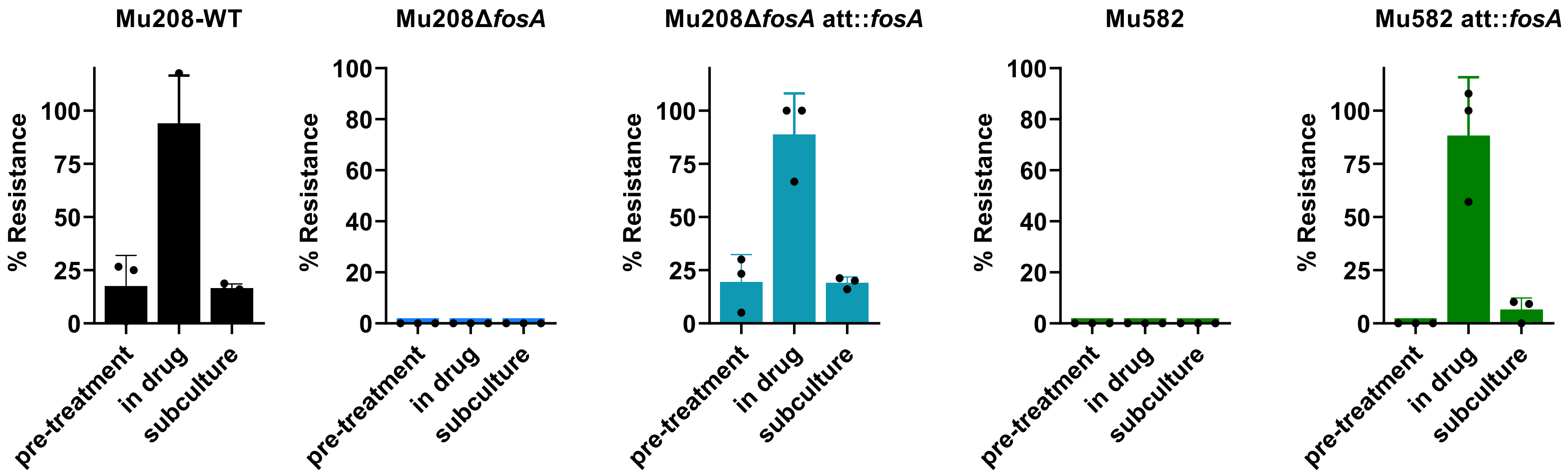


**Supplemental Figure 1. Fosfomycin resistant subpopulations in heteroresistant isolates are phenotypically unstable.** The frequency of the fosfomycin-resistant subpopulation of the indicated strains was quantified after growth in MH broth with G6P overnight (Pre-treatment), subsequent passage in MH broth with G6P and 128 µg/mL fosfomycin (in drug) for 24 h and subsequently in MH broth with G6P and no fosfomycin for 24 hours (subculture). The % resistance was calculated compared to the CFUs on no antibiotic plates.

Supplemental Figure 2

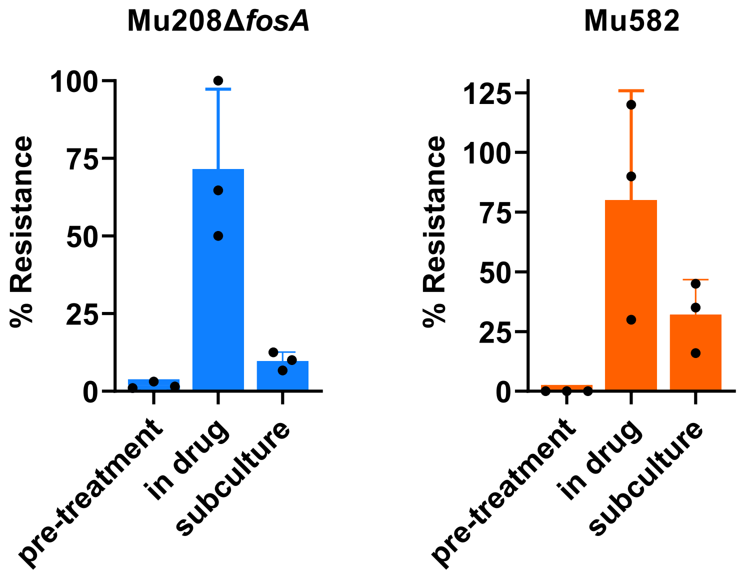


**Supplemental Figure 2. Heterosusceptibility is characterized by unstable, phenotypic heterogeneity.** The frequency of the subpopulation of the indicated strains with increased MIC was quantified after growth in MH broth with G6P overnight (Pre-treatment), subsequent passage in MH broth with G6P and fosfomycin ("in drug"; 4 µg/ml for Mu208Δ*fosA*, 1 µg/ml for Mu582) for 24 h and following passage in MH broth with G6P and no fosfomycin for 24 hours (subculture). The % resistance was calculated compared to the CFUs on antibiotic-free plates.

Supplemental Figure 3

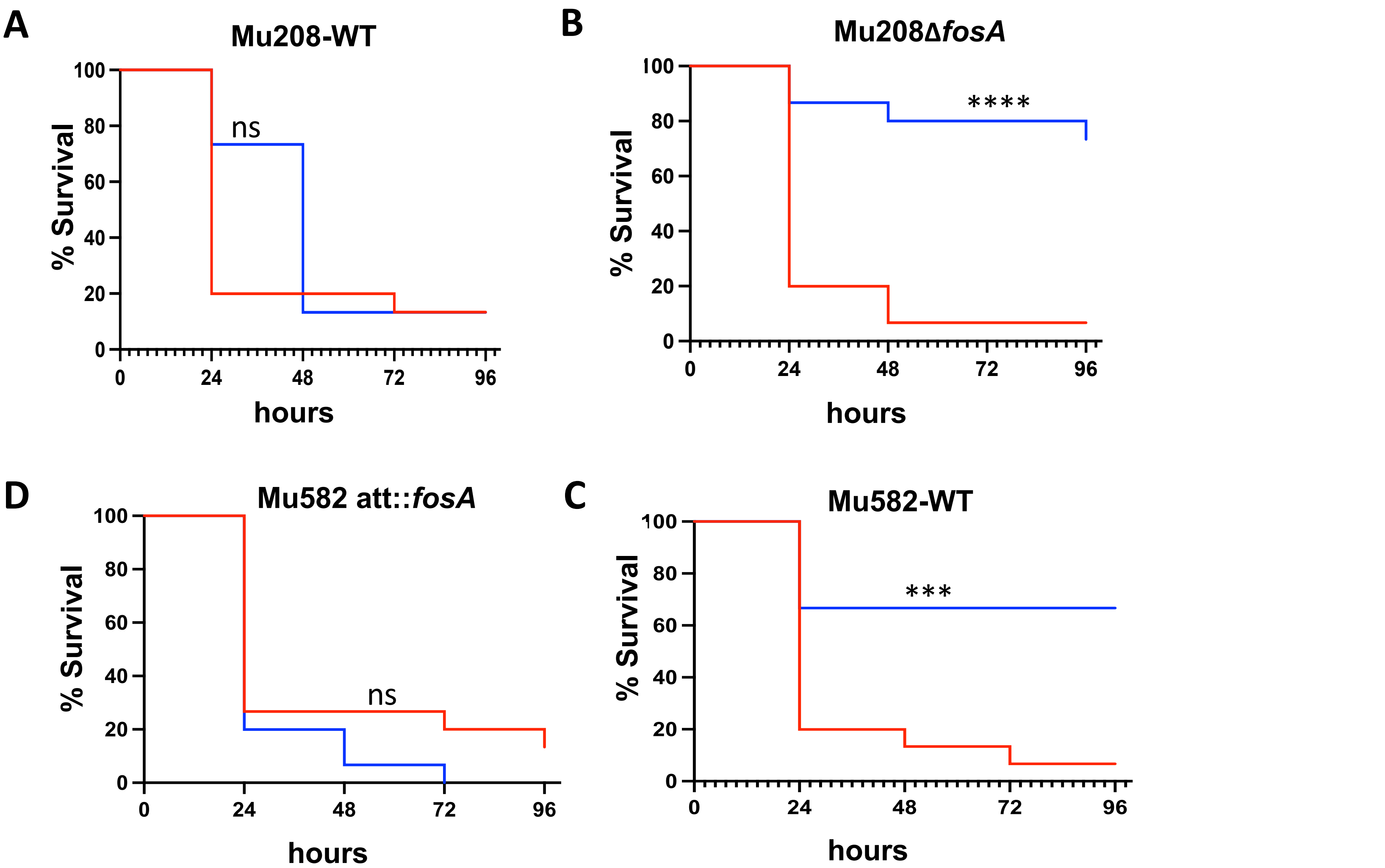


**Supplemental Figure 3. Heterosusceptibility does not mediate fosfomycin treatment failure.** *Galleria mellonella* (n = 15 per group) were infected with (A) Mu208-WT, (B) Mu208 Δ*fosA*, (C) Mu582-WT or (D) Mu582 att::*fosA* strains and then inoculated with PBS (untreated, red) or fosfomycin (treated, blue) 2 hours post-infection. Survival was monitored for 96 hours with data recorded every 24 hours. ****P* <0.001 and *****P* <0.0001, two-sided log-rank test.

Supplemental Figure 4

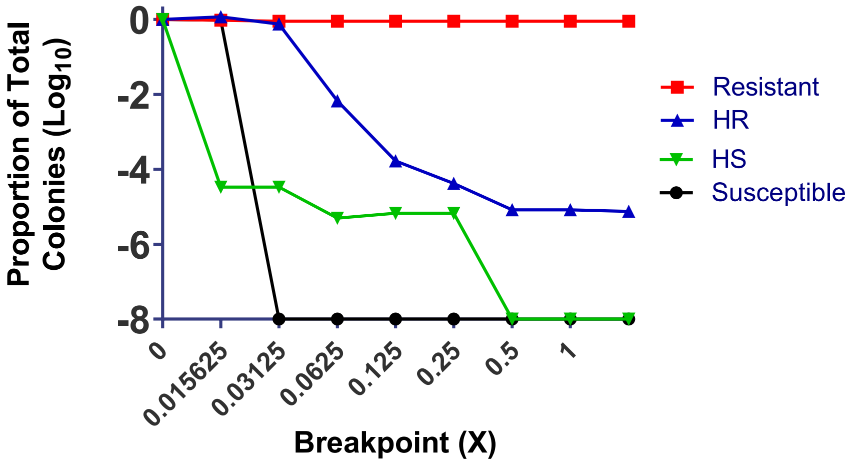


**Supplemental Figure 4. Schematic of population analysis profile (PAP) classifications for distinct resistance phenotypes.** The red squares represent data from a resistant isolate, in which all the cells survive at the breakpoint concentration. Blue triangles represent data from a heteroresistant isolate, in which there is a subpopulation of cells that survive at and beyond the breakpoint. Green inverted triangles represent data from a heterosusceptible isolate, in which there is a subpopulation of cells that are killed by an antibiotic concentration lower than the breakpoint. Black circles represent data from a susceptible isolate that does not contain a resistant subpopulation and is uniformly killed off at a low antibiotic concentration.
